# Analyzing matched sets of microbiome data using the LDM and PERMANOVA

**DOI:** 10.1101/2020.03.06.980367

**Authors:** Zhengyi Zhu, Glen A. Satten, Caroline Mitchell, Yi-Juan Hu

## Abstract

**Background:** Matched-set data arise frequently in microbiome studies. For example, we may collect pre- and post-treatment samples from a set of individuals, or use important confounding variables to match data from case participants to one or more control participants. Thus, there is a need for statistical methods for data comprised of matched sets, to test hypotheses against traits of interest (e.g., clinical outcomes or environmental factors) at the community level and/or the OTU (operational taxonomic unit) level. Optimally, these methods should accommodate complex data such as those with unequal sample sizes cross sets, confounders varying within sets, as well as continuous traits of interest.

**Methods:** PERMANOVA is a commonly used distance-based method for testing hypotheses at the community level. We have also developed the linear decomposition model (LDM) that unifies the community-level and OTU-level tests into one framework. Here we present a strategy that can be used with both PERMANOVA and the LDM for analyzing matched-set data. We propose to include an indicator variable for each set as covariates, so as to constrain comparisons between samples within a set, and also permute traits within each set, which can account for exchangeable sample correlations. The flexible nature of PERMANOVA and the LDM allows discrete or continuous traits or interactions to be tested, within-set confounders to be adjusted, and unbalanced data to be fully exploited.

**Results:** Our simulations indicate that our proposed strategy outperformed alternative strategies in a wide range of scenarios. Using simulation, we also explored optimal designs for matched-set studies. The flexibility of PERMANOVA and the LDM for a variety of matched-set microbiome data is illustrated by the analysis of data from two real studies.

**Conclusions:** Including set indicator variables and permuting within sets when analyzing matched-set data with PERMANOVA or the LDM is a strategy that performs well and is capable of handling the complex data structures that frequently occur in microbiome studies.

## Background

Many studies of the microbiome have matched-pair or matched-set designs, in which data naturally cluster into sets but the samples within each set vary in the traits of interest (e.g., clinical outcomes, environmental factors). Matching allows us to study the association between the microbiome and the traits of interest by comparing samples *within* sets, ignoring the variability in microbiomes *between* sets. For example, we may collected paired samples pre and post treatment from a set of subjects to assess the treatment effects on the microbiome. We may also collect matched case-control subjects who were matched on important confounding factors to facilitate case-control comparison. Matching is advantageous when the signal-to-noise ratio is larger within than between sets. In matched studies, complexities may occur when the data are *unbalanced* (e.g., having unequal ratio of case-to-control samples per set), there exist additional confounders that vary within each set, or some traits of interest are continuous.

Some methods have been developed specifically for modeling matched-set microbiome data but are limited to paired data without any within-pair covariates. Shi and Li [1] proposed a paired-multinomial distribution that is only applicable when the sample size is larger than the number of taxa. Zhao et al. [2] developed a generalized paired Hotelling’s test that relaxed the restriction of Shi and Li’s method, but can only provide tests at the community level. Some strategies have also been proposed to extend existing tests of the microbiome to analyzing matched-set data. The manual for the DESeq2 software package [3] stated that the set indicators should be included as a term in the design formula, but DESeq2 does not account for within-set correlations. The documentation of both implementations of PERMANOVA [4], `adonis2` (R package `vegan`) and `permanovaFL` (R package `ldm` [5]), suggest that restricted permutation within each set should be performed. (Note that the two implementations differ slightly in their permutation schemes; in particular, `permanovaFL` was based on the Freedman-Lane scheme [6]). However, the performance of these strategies have not yet been evaluated, especially in studies with unbalanced data or within-set confounders.

We previously introduced the linear decomposition model (LDM) [5] primarily for analyzing independent data. The LDM provides tests at the community level and also tests of the contribution of individual operational taxonomic units (OTUs; here we use “OTU” generically to refer to any feature such as amplicon sequence variants or taxonomic/functional grouping of bacterial sequences). These tests are conducted in a unified manner such that the findings of a community-level test can be resolved with the findings at the individual OTU level. Both PERMANOVA and the LDM are regression- and permutation-based, making them readily extendable to analyzing matched-set data while accounting for the aforementioned data complexities. Although we considered within-cluster permutation in the LDM paper [5], that was in a context in which variables of interest could be below the cluster level. We did not explicitly consider the matched set data we describe here from either a theoretical or numerical point of view.

In this article, we develop a new strategy for using PERMANOVA and the LDM to analyze a wide range of matched-set microbiome data. In the methods section, we describe our strategy and establish a connection with the existing strategy of restricted permutation. In the results section, we present the simulation studies and the application to two real microbiome studies with matched-set designs. We conclude with a discussion section.

## Methods

We will refer to each observation as a “sample” and refer to the experimental unit that contributes one or more observations as a “set”. We allow each set to be comprised of an arbitrary number of samples. We also allow multiple discrete and/or continuous traits to be tested and additional sample-level (i.e., within-set) confounding covariates to be adjusted for. In a common scenario with a binary trait (e.g., a case-control status or a treatment or exposure variable), each set consists of one case sample and *m* (*m* ≥ 1) control samples, usually referred to as 1:*m* matched data. We assume that, after all covariates (including the traits of interest) have been accounted for, the members of each set are exchangeable.

To present our strategy for analyzing matched-set data, we introduce a common notation to describe both PERMANOVA and the LDM. Both PERMANOVA and the LDM are linear models for which the effects of covariates (metadata) are summarized in a design matrix *X*. The rows of *X* correspond to samples while the columns of *X* correspond to the covariates. We may partition *X* by columns into *K* groups (which we call “submodels”) such that *X* = (*X*_1_, *X*_2_, …, *X*_*K*_), where each *X*_*k*_ denotes a variable or set of variables we wish to test (jointly). For example, *X*_*k*_ may consist of indicator variables for levels of a single categorical variable, or a group of potential confounders that we wish to adjust for simultaneously. Both PERMANOVA and the LDM make the columns of *X*_*k*_ *orthonormal* to the columns of *X*_*k′*_ for *k*′ *< k* using projection (i.e., the Gram-Schmidt process).

For both PERMANOVA and the LDM, test statistics for the *k*th submodel can be expressed in terms of the quantity 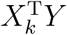. For PERMANOVA, *Y* is related to the (squared and Gower-centered) distance matrix Δ by Δ = *Y SY* ^T^, where *S* is a diagonal matrix with diagonal elements equal to 1 or −1 corresponding to positive and negative eigenvalues, respectively. For the LDM, *Y* is the (column-centered) OTU table that has rows for samples and columns for OTUs; the OTU table typically contains the frequency (i.e., relative abundance) data or arcsin-root transformed frequency data. Since *Y* in either PERMANOVA or the LDM is column-centered and treated as the response of a linear model, we also assume the design matrix *X* is *column-centered*.

With no loss of generality, we can write the element of *Y* in the *i*th row and *j*th column as

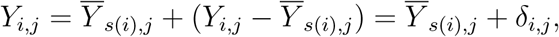

where *s*(*i*) is the set that the *i*th sample belongs to. Thus 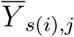 is the set-level average of *Y*_*i,j*_ and *δ*_*i,j*_ is the deviation of the *i*th sample from the set-level average. The rationale of a matched-set design is that we wish to treat 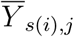 characterizing a set as a *nuisance parameter* and focus the testing efforts on *δ*_*i,j*_s. With this in mind, we note that 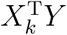 is a function of only the *δ*_*i,j*_s (i.e., not a function of the 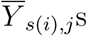) whenever the column values of *X*_*k*_ sum to zero for each set of samples belonging to the same set. It is clear that this occurs whenever the columns of *X*_*k*_ are orthogonal to the set of indicator variables corresponding to the set IDs. Therefore, our proposed strategy for fitting matched-set data is to introduce an indicator variable for each set to be included in submodel *X*_1_ along with any sample-level confounding covariates that are not matched on. Note that any set-level confounders are automatically controlled for in this strategy, as they can be written as linear combinations of the indicator variables generated by the set IDs. Indeed, it is typical of matched-set analyses that the effect of variables that have been matched on (i.e., that are constant in each set) cannot be determined (see e.g., [7]).

To see how this works in practice, consider a simple example with two sets, the first having two samples and the second having three samples. For clarity, we work with *X*_*k*_s before orthonormalization and show *X*_1_ (which has the indicator variables for the two sets) and *X*_2_ (which has a case-control status) before column centering:

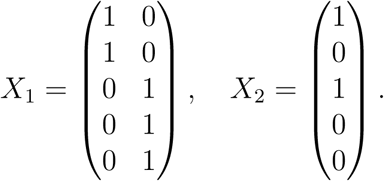

After column centering (i.e., subtracting column means), *X*_2_ = (3*/*5, −2*/*5, 3*/*5, −2*/*5, −2*/*5)^T^. Note that the values in *X*_2_ do not sum to zero within each set. If we constructed a test using this contrast, the set-specific means 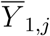 and 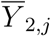 would not be eliminated. However, if we make *X*_2_ (before column centering) orthogonal to the columns of *X*_1_ (which automatically achieves column centering), we find *X*_2_ = (1*/*2, −1*/*2, 2*/*3, −1*/*3, −1*/*3)^T^, where we see that the values in *X*_2_ sum to zero within each set.

We identify a condition under which the nuisance parameters disappear even without projecting off the set ID. We say that a variable in matched-set data is *balanced* if the sum of the variable within each set is proportional to the number of its samples (with the same constant of proportionality). For example, a case-control status is balanced if all sets have as many case as control samples, or if some sets have two case and four control samples and the remaining sets have one case and two control samples. *For a balanced variable, column centering alone is sufficient to make the values of that variable sum to zero within each set, even without projecting off the set ID*. Note that adjusting for sample-level covariates can result in imbalance in a variable, even if it was initially balanced; in this case, projection on the set ID is required to restore balance. A simple example with two sets, each contributing two samples along with a sample-level covariate, shows this. Before column centering (and orthonormalization), suppose the covariate is *X*_1_ = (9, 8, 6, 9)^T^ and the case-control status is *X*_2_ = (1, 0, 1, 0)^T^. After column centering we have

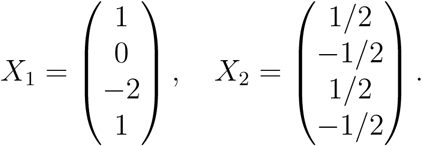

In the absence of the covariate, *X*_2_ after column centering do sum to zero within each set; however, after adjusting for the covariate we have *X*_2_ = (2*/*3, −1*/*2, 1*/*6, −1*/*3)^T^, which does not eliminate the set-specific means. If we also adjust for the set ID by augmenting *X*_1_ with the column-centered indicator

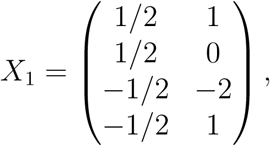

we obtain *X*_2_ = (3*/*5, −3*/*5, 1*/*5, −1*/*5)^T^, in which values do sum to zero within sets. Finally, note that in this example we have considered a binary case-control trait; it should be clear that, for a continuous trait, the within-set sum is unlikely to be the same for each set, and hence the projection on the set ID is required to eliminate the nuisance parameters.

As we have assumed the samples in each set are exchangeable, we propose to perform restricted permutation among samples from the same set. As permuting residuals of *Y* in the Freedman and Lane scheme [6] is typically equivalent to permuting *X*_*k*_s [5], the restricted permutation refers to permuting the (orthonormalized) traits of interest within samples from the same set. The same permutation scheme can be used for testing the interactions between traits of interest or between traits and set-level covariates; the latter allows us to detect whether the associations between the microbiome and the traits of interest are homogeneous across study groups (which can be defined by the set-level covariates). As noted previously, when *all* variables are balanced, the columns of *X* (excluding the set ID indicator vectors) will automatically be orthogonal to the set ID indicator vectors. Since both `permanovaFL` and the LDM permute the rows of *X*, it is also clear that this orthogonality holds for every permutation as long as permutations are conducted within sets. As a result, the *p*-values for `permanovaFL` or the LDM will be identical with and without adjustment of the set ID in this situation, as long as the restricted permutation is performed.

## Results

### Simulation studies

To generate our simulation data, we used the same motivating dataset as Hu and Satten [5], i.e., data on 856 OTUs of the upper-respiratory-tract microbiome first described by Charlson et al. [8]. In most simulations we considered a binary trait such as case-control status, but we also considered matched sets with a continuous trait. We defined a “causal” OTU to have frequency that depended on the trait of interest. We considered on two complementary causal mechanisms: the first mechanism (S1) assumed that half (428) of the OTUs (after excluding the three most abundant OTUs) were causal; the second mechanism (S2) assumed the ten most abundant OTUs were causal. In each scenario, we randomly partitioned the causal OTUs into two equal-size subsets, 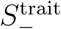 and 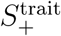, to contain OTUs with decreased and increased frequencies, respectively, in cases relative to controls. We further partitioned 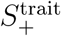 into 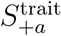 and 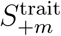 comprised of OTUs whose frequencies are increased in additive and multiplicative manners, respectively. For the simple situation, with no covariates but the trait of interest, we simulated data for the *i*th set using the following steps.

1. We assigned trait values 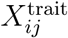 to the *j*th sample of the *i*th set. For matched pair samples, 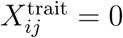 was assigned to control samples and 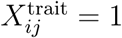 to case samples; for continuous traits, 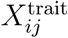 was sampled from the *U* [0, 1] distribution.
2. We generated the mean OTU frequencies 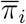 for set *i* from the Dirichlet distribution 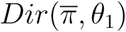, where the mean parameter 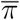 and overdispersion parameter *θ*_1_ took the values of the estimated mean and overdispersion (0.02) in the Dirichlet-Multinomial (DM) model fitted to the upper-respiratory-tract data. Note that 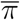 and *θ*_1_ characterize the population mean of OTU frequencies and between-set heterogeneity.
3. Given 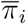, we generated the baseline OTU frequencies 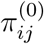 for sample *j* of set *i* from the Dirichlet distribution 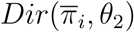, where *θ*_2_ was set to 0.007, which was the median of the estimated overdispersion in the DM model that was fitted to data for each set with three samples in the MsFLASH study (see the section “Analysis of the MsFLASH data”). Note that *θ*_2_ characterizes heterogeneity among samples from the same set, and 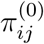 represents the (true) OTU frequencies we would see when trait 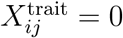.
4. We then generated the (true) OTU frequencies that account for a non-zero effect of trait 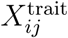, denoted 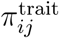, by reducing the frequency of each OTU in 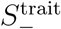 by a factor of *β*, then distributing half of the total reduced frequency evenly to OTUs in 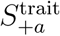 and the other half to OTUs in 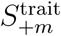 in proportion to their baseline frequencies in 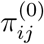. We then formed the (true) OTU frequency for the *j*th sample from the *i*th set using 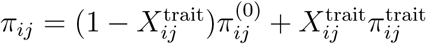. Note that *β* characterizes the *effect size* of the trait, i.e. the amount by which OTU frequencies vary at the causal OTUs when the trait 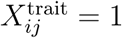.
5. We generated read count data for each sample using the multinomial distribution *MN* (*π*_*ij*_, *N*_*ij*_), where the total read count *N*_*ij*_ was generated from the Poisson distribution with mean 10000 and left truncated at 500.

To induce the effects of additional covariates, we made further modifications to 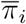 and/or *π*_*ij*_ that were similar to the modifications made to 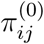 to construct 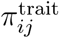. For simulations where we wished to include a main effect of a set-level covariate 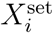, we first sampled values of 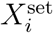 from a Bernoulli distribution with parameter 0.5. We then uniformly sampled 428 OTUs to be associated with the covariate, and randomly partitioned them into two equal-size subsets 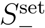 and 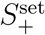. We then constructed 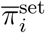 by modifying 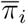, reducing the frequency of each OTU in 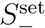 by a factor of *β*^set^ = 0.2 and distributing the total reduced frequency to OTUs in 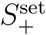 in proportion to their original frequencies in 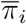. We then replaced 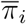 by 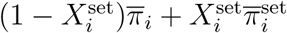, to be used in Step 3.

To account for a sample-level confounder 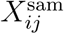, we first sampled 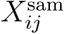 from a Bernoulli distribution with parameter 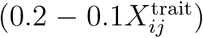. We then uniformly sampled 428 OTUs to be associated with the covariate, and randomly partitioned them into two equal-size subsets 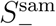 and 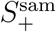. We then constructed 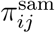 by modifying *π*_*ij*_ in the same way that 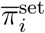 was modified from 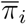, but with a factor of *β*^sam^ = 0.5. We then replaced *π*_*ij*_ by 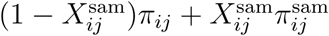. The resulting values were used in Step 5.

Finally, to account for an interaction between a set-level covariate and the trait, we sampled a third set of OTUs (a random sample of 428 OTUs under S1 and the top 1-5 and 11-15 most abundant OTUs under S2) to be associated with the interaction, and randomly partitioned them into two equal-size subsets 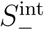 and 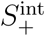. Then, when both 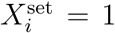 and 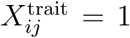, we further modified *π*_*ij*_ by reducing the frequency of OTUs in 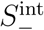 by a factor *β*^int^ and then distributing this extra mass to 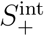 in proportion to the OTU frequencies in *π*_*ij*_. The resulting values of *π*_*ij*_ were then used in Step 5. Note that whenever we included an interaction term like this, the main effect of 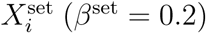 and 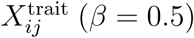 was also included as described previously.

We evaluated the performance of different strategies and methods in seven scenarios of matched-set data: (1) matched-pair data, (2) unbalanced data, (3) matched-pair data with a sample-level confounder, (4) matched-pair data with a set-level covariate, (5) unbalanced data with a set-level covariate, (6) matched-pair data with a continuous trait, and (7) matched-pair data with an interaction effect. To facilitate comparison across scenarios, the same sets of causal OTUs 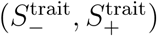 and covariate-associated OTUs 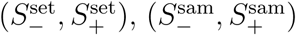 and 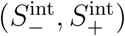 (if called for), were used for all scenarios. For each scenario except (2), (5) and (6), we generated data for 50 1:1 matched pairs (with a binary trait); for scenarios (2) and (5) with unbalanced data we generated data for 25 1:1 matched pairs and 25 1:2 matched sets (with a binary trait); for scenario (6), we generated data for 50 matched pairs with a continuous trait.

We also explored various 1:*m* matched study designs to assess the performance under varying conditions. First, we compared the design that collected 50 1:1 matched pairs with the design that collected 50:50 independent case-control samples (first simulating pairs and then selecting only one sample from each pair), over varying values for the within-set heterogeneity *θ*_2_. Second, we compared different 1:*m* matched-set designs with a fixed total of 90 samples. Specifically, we considered *m* = 1, 2, 4, and 5 and collected 45 1:1 pairs, 30 1:2 sets, 18 1:4 sets, and 15 1:5 sets, respectively, to form each dataset. We also considered *m* = 3 and collected 22 of 1:3 sets and 1 pair (to meet the total sample size 90) for the 1:3 design. Lastly, we compared different 1:*m* (*m* = 1, 2, 3, 4, 5) designs when fixing the total number of sets to 50.

We applied PERMANOVA (implemented in both `permanovaFL` and `adonis2`) and the LDM with the proposed strategy (*adjusting for the set ID and sample-level covariates if present, not adjusting for set-level covariates, and performing restricted permutation within sets*). PERMANOVA tests were calculated using the Bray-Curtis distance unless otherwise noted. We report LDM results for the omnibus test that combines the test results from raw frequency (relative abundance) data and arcsin-root-transformed frequency data [5]. We compared the proposed strategy to alternative strategies, specifically: not adjusting for the set ID; or not performing restricted permutation. We compared our results to DESeq2 in testing individual OTUs, which adjusts for the set ID but does not account for sample correlations. We also compared our results to the Wilcoxon signed-rank test when it is applicable (i.e., for matched-pair data). We evaluated the type I error and power for the community-level (global) test of any microbiome effect at nominal significance level 0.05, and we assessed empirical sensitivity (proportion of truly associated OTUs that were detected) and empirical FDR for the OTU tests at a nominal FDR of 10%. Results for type I error were based on 10000 replicates; all other results were based on 1000 replicates. OTUs having fewer than 5 non-zero entries were removed before analysis.

### Simulation results

Results on type I error for the seven scenarios we considered were summarized in Table 1. The results of power, sensitivity, and FDR for the seven scenarios were displayed in Figures 1–7, respectively. In all scenarios, our proposed strategy, when applied to either `permanovaFL` or the LDM, yielded correct type I error and the highest power compared to alternative strategies; `adonis2` with the proposed strategy produced slightly conservative type I error and slightly lower power compared to `permanovaFL`. The LDM using the proposed strategy always controlled the FDR and achieved the highest sensitivity compared to the LDM using alternative strategies or DESeq2 when it controlled the FDR or Wilcoxon if it is applicable. In what follows, we describe the performance of these methods using other alternative strategies in each scenario.

**Table 1.** Type I error for testing the community-level hypothesis at level 0.05.

| Scenario and Analysis Strategy | permanovaFL | adonis2 | LDM |
| --- | --- | --- | --- |
| (1) Matched-pair data |  |  |  |
| Proposed | 0.0491 | 0.0450 | 0.0479 |
| Not adjusting for ID | 0.0491 | 0.0441 | 0.0479 |
| Unrestricted permutation | 0.0024 | 0.0855 | 0.0169 |
| (2) Unbalanced data |  |  |  |
| Proposed | 0.0471 | 0.0434 | 0.0505 |
| Not adjusting for ID | 0.0501 | 0.0456 | 0.0500 |
| Unrestricted permutation | 0.0039 | 0.0732 | 0.0280 |
| (3) Matched-pair data with a sample-level confounder $X^{\text{sam}}$ | | | |
| Proposed | 0.0452 | 0.0429 | 0.0476 |
| Not adjusting for $X^{\text{sam}}$ | 0.0688 | 0.0631 | 0.0800 |
| Not adjusting for ID | 0.0713 | 0.0328 | 0.0872 |
| Unrestricted permutation | 0.0016 | 0.0810 | 0.0163 |
| (4) Matched-pair data with a set-level covariate $X^{\text{set}}$ | | | |
| Proposed | 0.0510 | 0.0433 | 0.0481 |
| Not adjusting for ID | 0.0510 | 0.0451 | 0.0481 |
| Not adjusting for ID, adjusting for $X^{\text{set}}$ | 0.0510 | 0.0450 | 0.0481 |
| Unrestricted permutation | 0.0021 | 0.0874 | 0.0162 |
| (5) Unbalanced data with a set-level covariate $X^{\text{set}}$ | | | |
| Proposed | 0.0479 | 0.0444 | 0.0480 |
| Not adjusting for ID | 0.0496 | 0.0446 | 0.0483 |
| Not adjusting for ID, adjusting for $X^{\text{set}}$ | 0.0489 | 0.0445 | 0.0489 |
| Unrestricted permutation | 0.0048 | 0.0736 | 0.0257 |
| (6) Matched-pair data with a continuous trait |  |  |  |
| Proposed | 0.0505 | 0.044 | 0.0461 |
| Not adjusting for ID | 0.0677 | 0.0612 | 0.0881 |
| Unrestricted permutation | 0.190 | 0.906 | 0.994 |
| (7) Matched-pair data with an interaction effect |  |  |  |
| Proposed | 0.0524 | 0.0295 | 0.0536 |
| Unrestricted permutation | 0 | 0.0977 | 0 |

For each of the seven scenarios, results for three or four analysis strategies are presented. First in each scenario is the “Proposed” strategy that adjusts for the set ID indicators and sample-level covariates (if present), does not adjust for set-level covariates (if present), and performs restricted permutation within sets. Each alternative strategy is described by its difference from the proposed strategy; for example, “Unrestricted permutation” maintains all the elements of the proposed strategy except for replacing the recommended within-set permutation with an unrestricted permutation.

**Fig 1.**
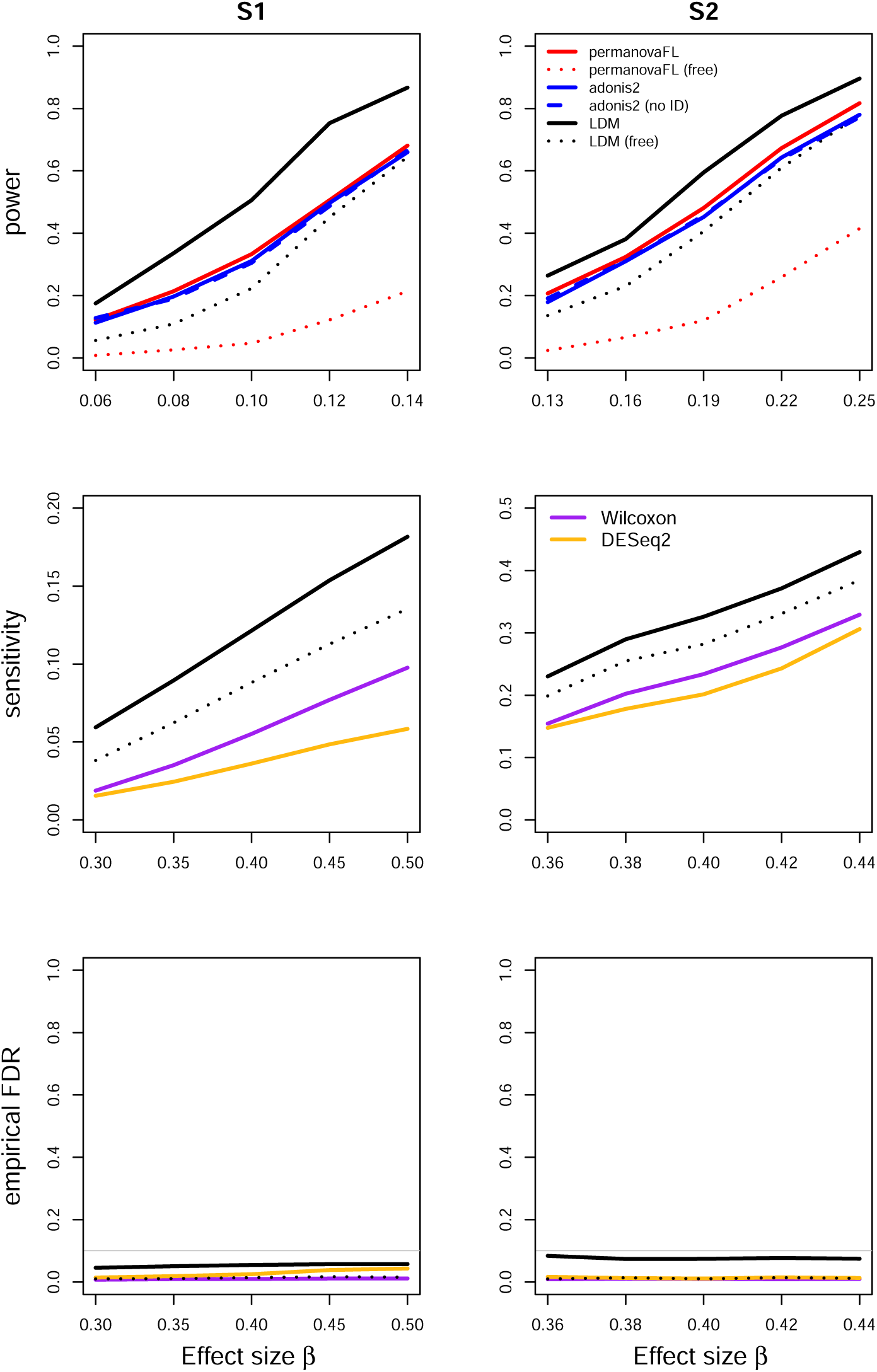
Simulation results for the matched-pair data of scenario (1). “Free” means unrestricted permutation; “no ID” means not adjusting for the set ID. Because the LDM and `permanovaFL` gave identical results with and without adjustment for set ID indicators, only results using the proposed strategy for these two methods are shown here. `adonis2` with unrestricted permutation had inflated type I error in all scenarios we examined and is therefore not shown in subsequent figures that display power or sensitivity.

For (1) the matched-pair data, `permanovaFL` and the LDM not adjusting for the set ID produced identical results to their counterparts using the proposed strategy as expected. Note that *p*-values from `adonis2` were not identical with and without adjustment for set ID, but the type I error and power (Figure 1) of the two strategies were very similar. Here and for all datasets with a binary case-control trait, the strategy of performing *unrestricted* permutation led to conservative type I error and FDR and diminished power and sensitivity (Figures 1–5, 7 when applied to `permanovaFL` and the LDM, but inflated type I error when applied to `adonis2`.

For (2) the unbalanced data, the LDM not adjusting for the set ID yielded correct type I error but diminished power and sensitivity relative to its counterpart that using the proposed strategy (Figure 2). The same pattern can be seen in the results of `permanovaFL` and `adonis2`.

**Fig 2.**
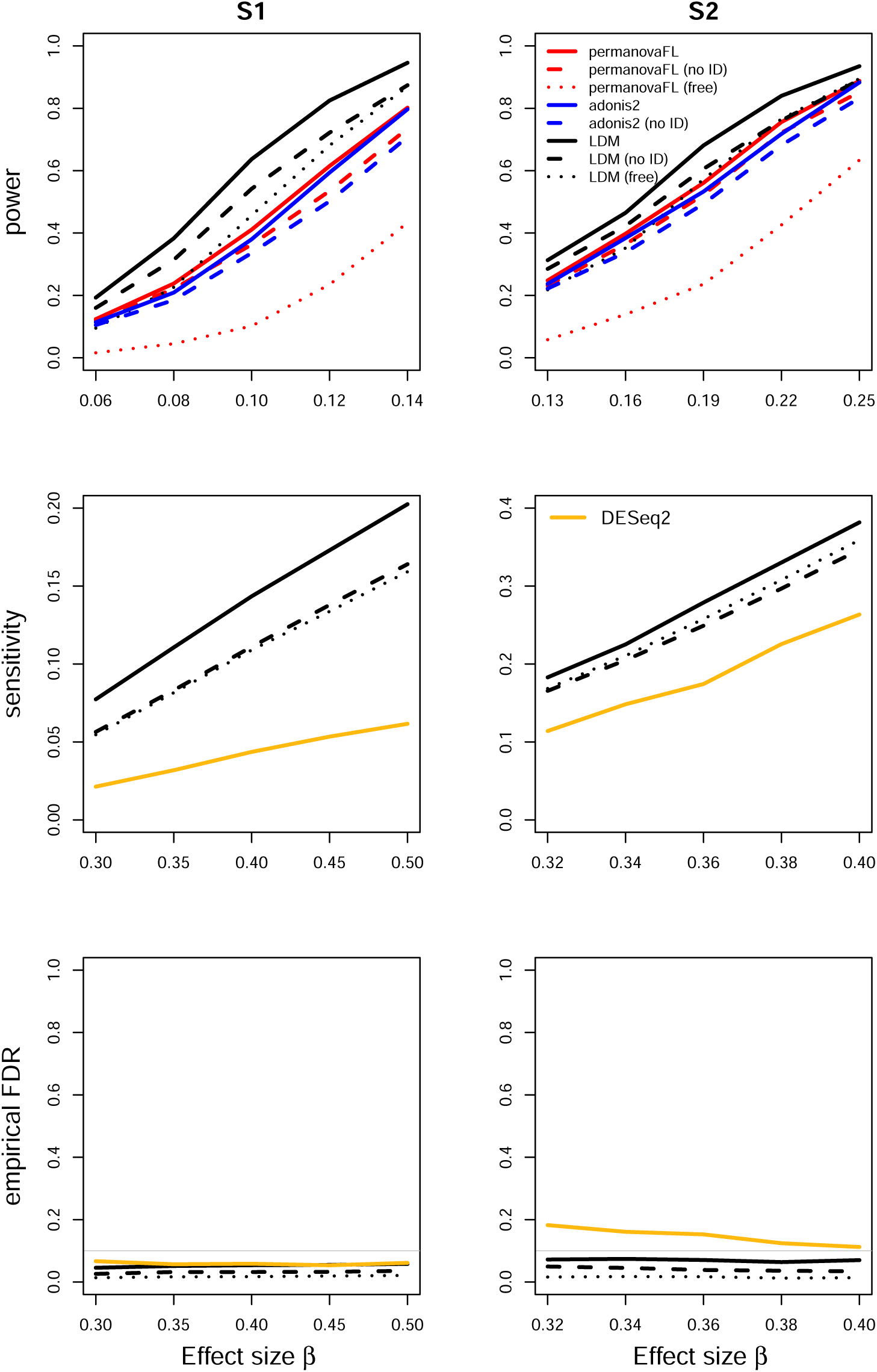
Simulation results for the unbalanced data of scenario (2).

**Fig 3.**
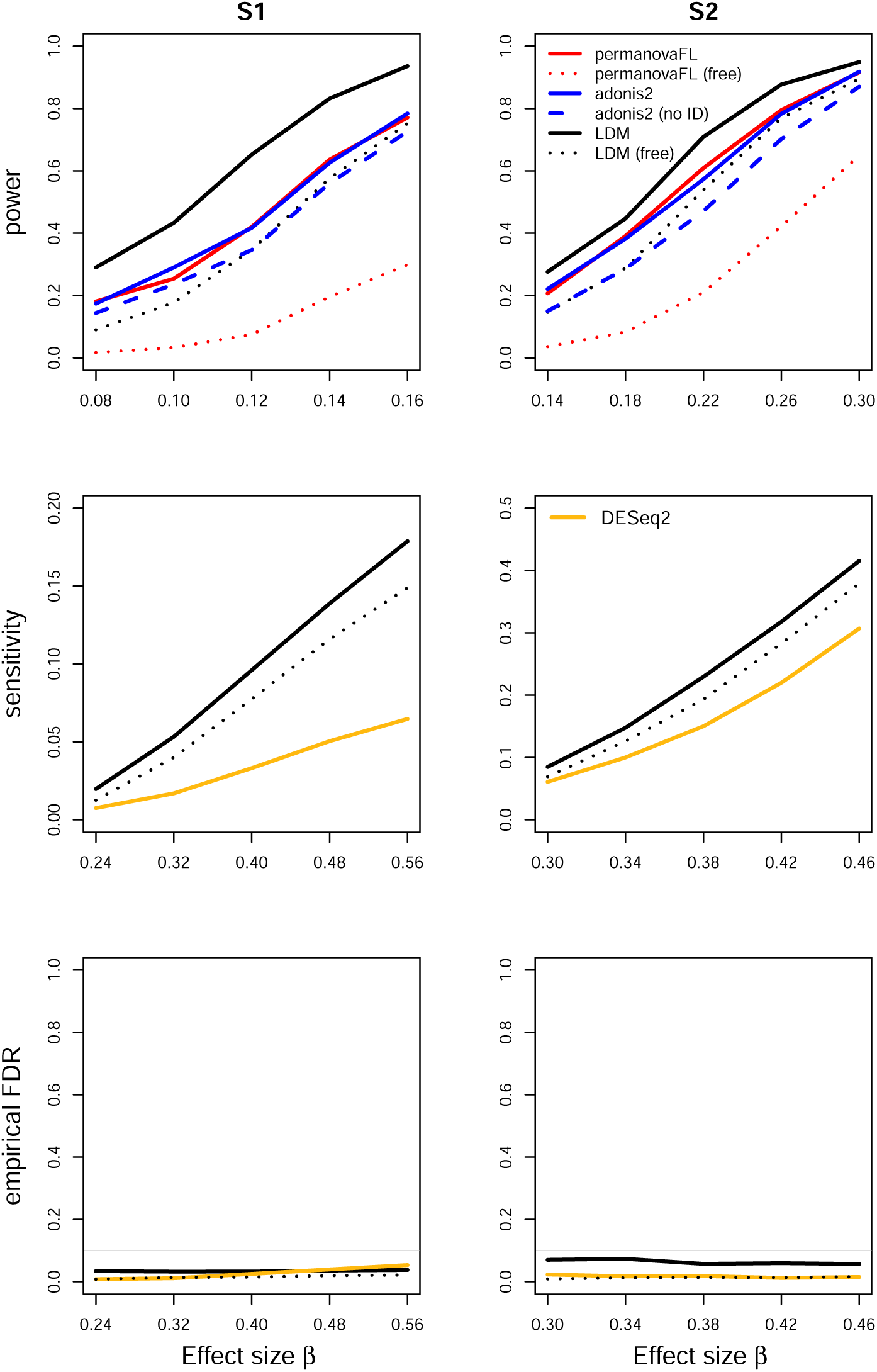
Simulation results for the matched-pair data with a sample-level confounding covariate of scenario (3). “no ID” means not adjusting for the set ID but adjusting for the confounder in this scenario. The LDM and `permanovaFL` with the “no ID” strategy had inflated type I error and thus are not shown.

**Fig 4.**
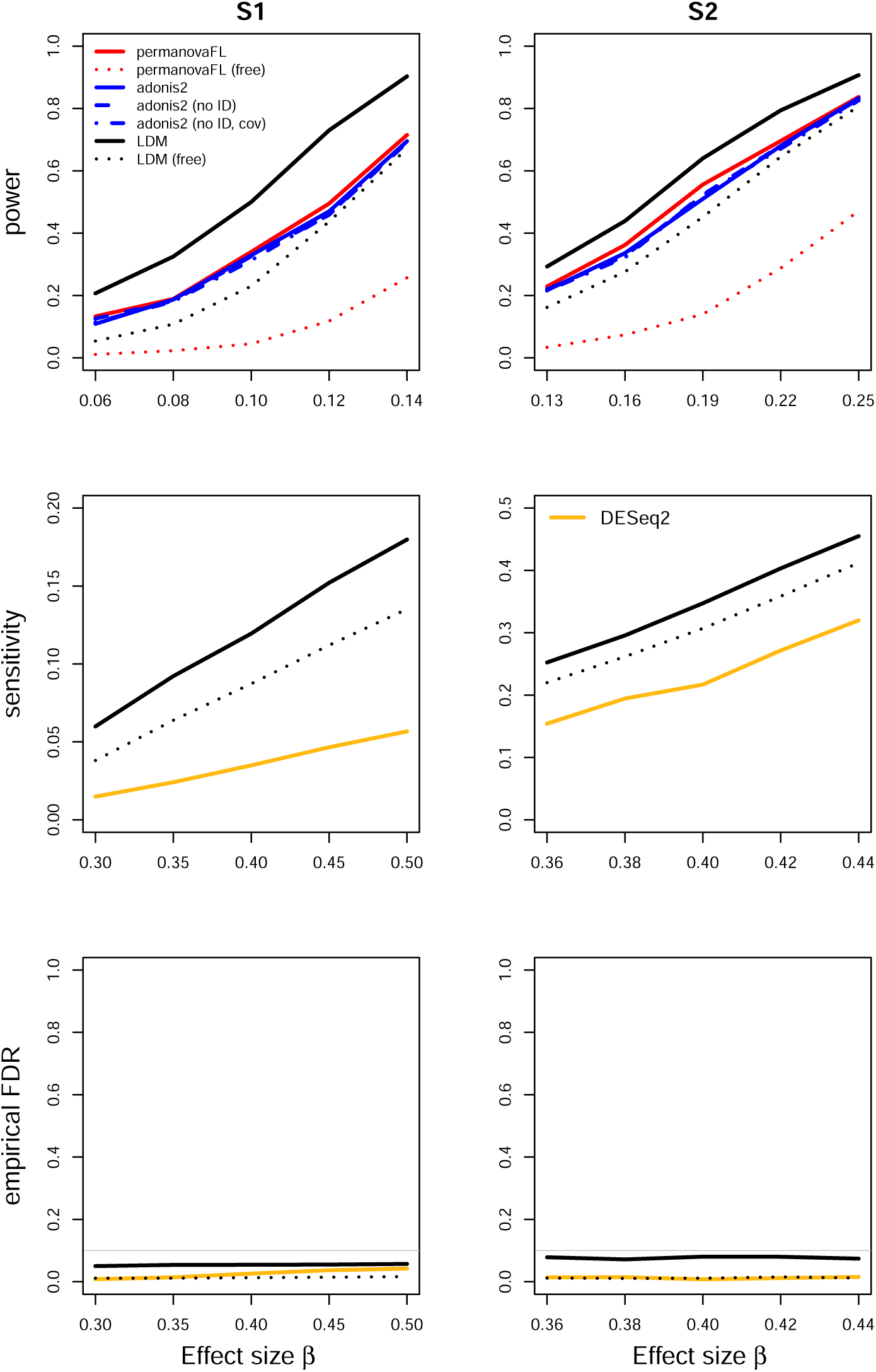
Simulation results for the matched-pair data with a set-level confounding covariate of scenario (4). “no ID” means not adjusting for the set ID (second strategy in scenario (4) of Table 1); “no ID, cov” means not adjusting for the set ID but adjusting for the covariate *X*^set^ (third strategy). The LDM and `permanovaFL` with “no ID” or “no ID, cov” had identical results as their counterparts with the proposed strategy and are thus not shown here.

**Fig 5.**
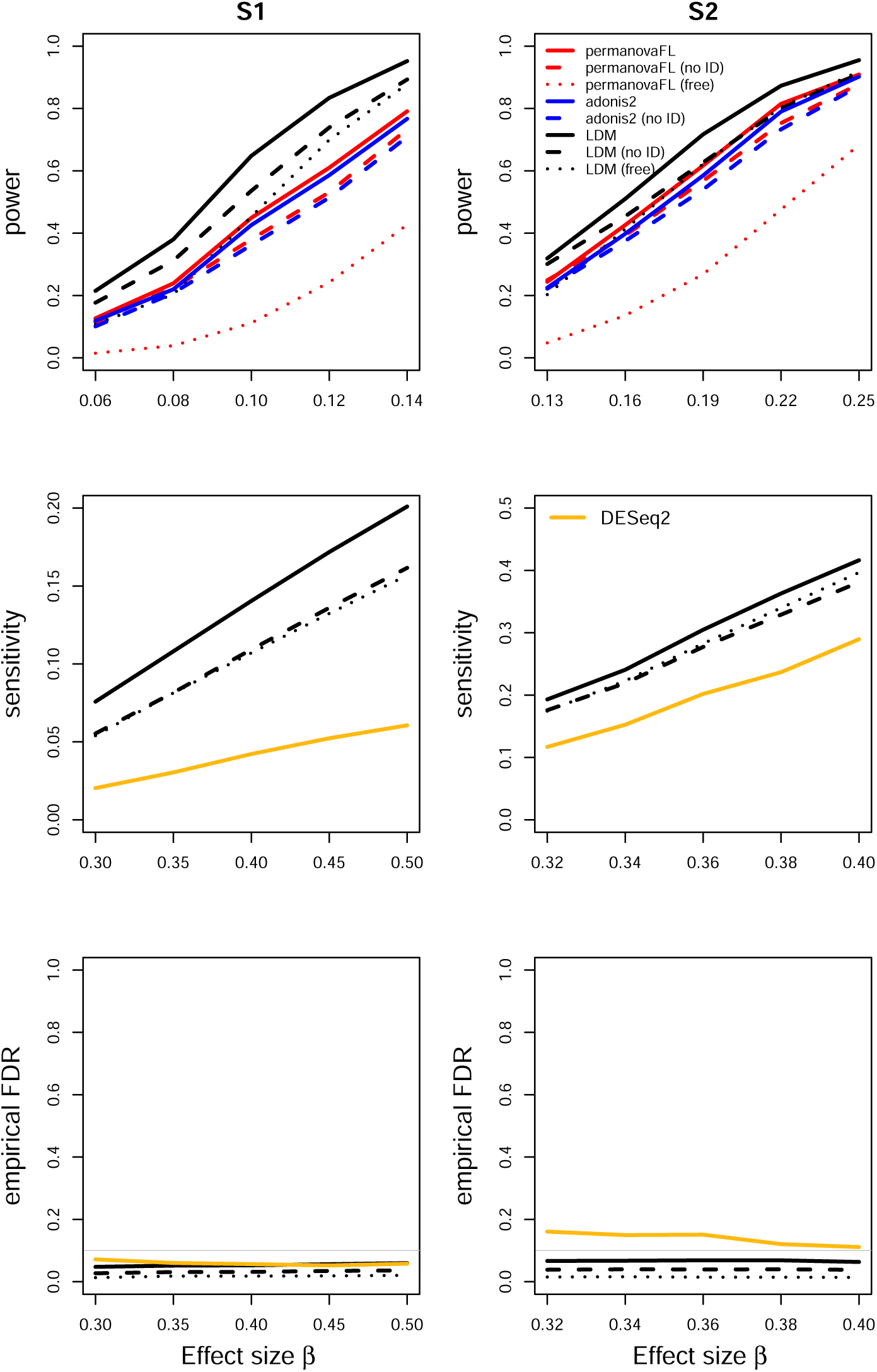
Simulation results for the unbalanced data with a set-level covariate of scenario (5). “no ID” means not adjusting for the set ID (second strategy in scenario (5) of Table 1). The power and sensitivity of methods using the strategy not adjusting for ID but adjusting for *X*^set^ (third strategy) are very similar to the power and sensitivity of their counterparts using the “no ID” strategy; thus only the latter is shown.

**Fig 6.**
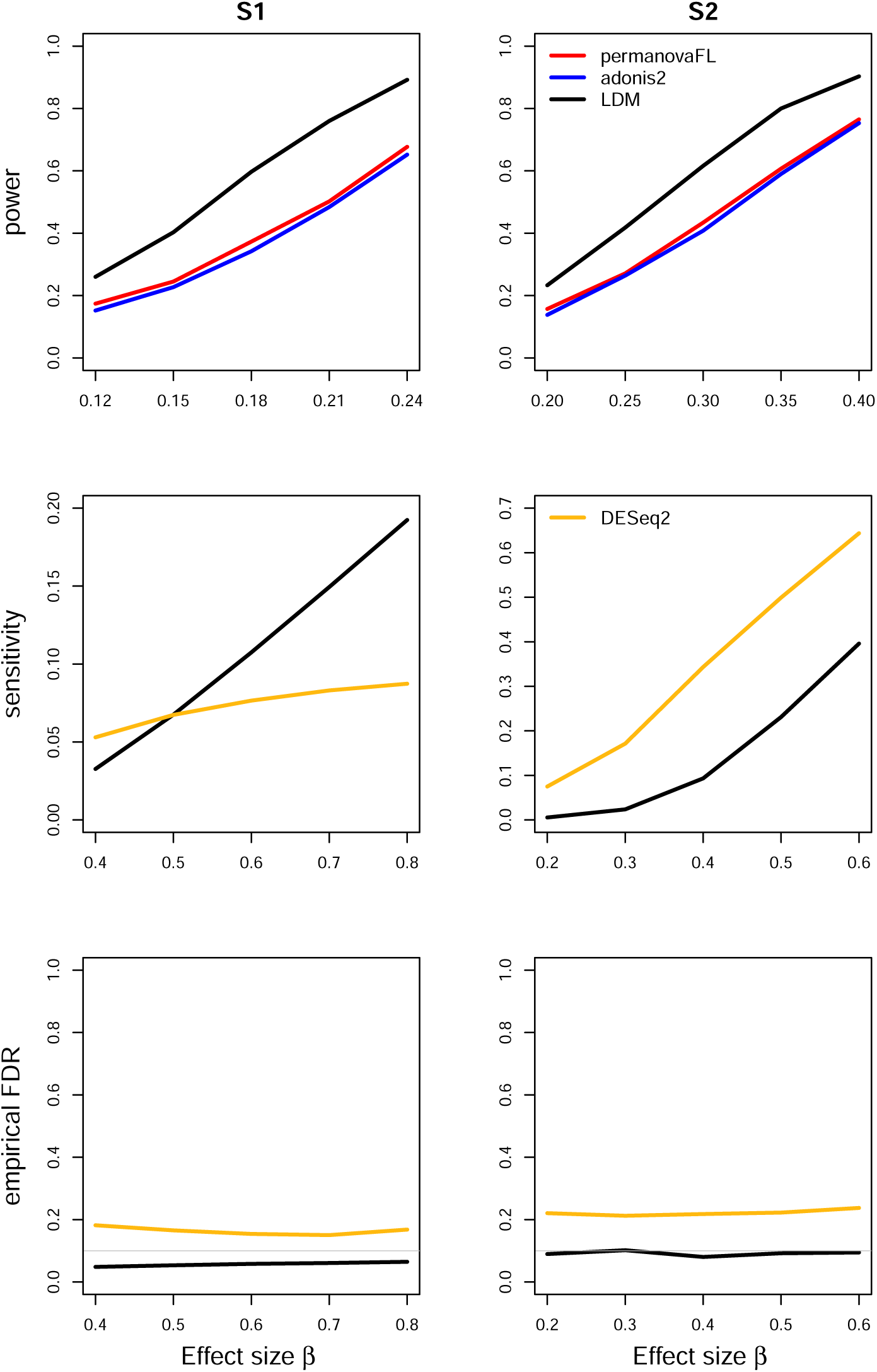
Simulation results for the matched-pair data with a continuous trait of scenario (6). The other strategies all led to inflated type I error, and are thus not shown here.

**Fig 7.**
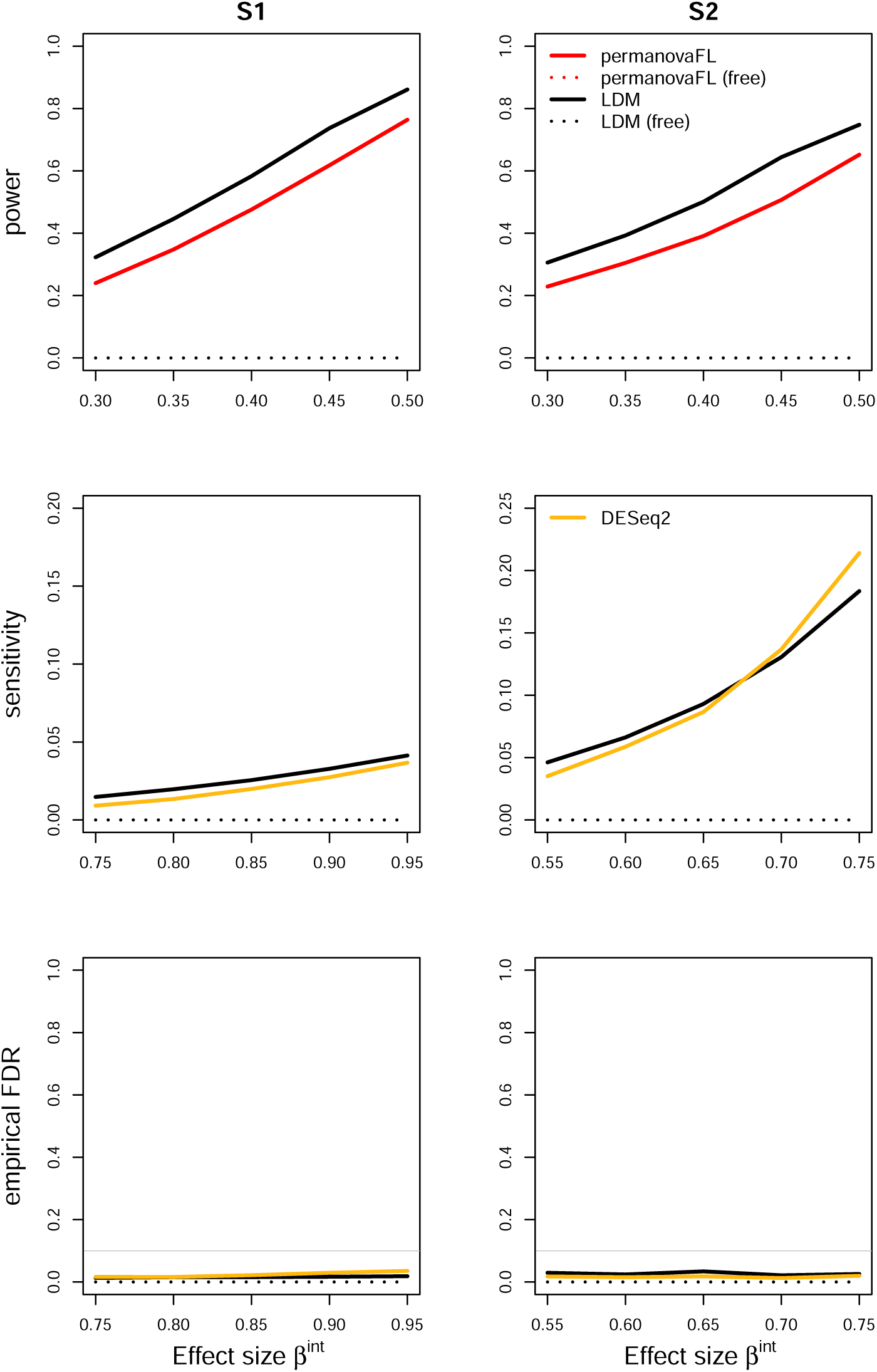
Simulation results for scenario (7) testing an interaction between a (set-level) group variable and a (sample-level) trait variable in matched-pair data.

For (3) the matched-pair data with a sample-level confounder, `permanovaFL`, `adonis2`, and the LDM not adjusting for the confounder had inflated type I error (0.063 ∼ 0.080), indicating that we have indeed induced some confounding effect in the data. In the presence of such a confounding effect, `permanovaFL` and the LDM not adjusting for the set ID (even after adjusting for the confounder) had inflated type I error (0.071 ∼ 0.087). In this case, not adjusting for the set ID did not just affect the power, but also affected the validity. These methods with inflated type I error were not included in Figure 3. In contrast, `adonis2` not adjusting for the set ID had more conservative type I error than that adjusting for the set ID (both adjusting for the confounder), so the former had reduced power compared to the latter.

Our proposed strategy was robust to the presence of set-level covariates. For (4) the matched-pair data with a set-level covariate, whether or not adjusting for the covariate or the set ID (i.e., the first three strategies in scenario (4) of Table 1) all yielded identical results when applied to `permanovaFL` or the LDM, as we have analytically shown. Thus Figure 4 only displayed their results for the proposed strategy. When applied to `adonis2`, the three strategies led to slightly different type I error and power. For (5) the unbalanced data with a set-level covariate, the LDM not adjusting for the set ID generated correct type I error but diminished power and sensitivity compared to its counterpart that adjusted for the set ID (both not adjusting for the covariate) (Figure 5). Adjusting for the covariate but not the set ID failed to recover any power or sensitivity, which understored the importance of adjusting for the set ID. The same pattern can be seen in the results of `permanovaFL` and `adonis2`.

In the presence of a continuous trait, even in (6) the simplest matched-pair data without any covariates, `permanovaFL`, `adonis2`, and the LDM not adjusting for the set ID all yielded inflated type I error. The strategy of performing unrestricted permutation led to highly inflated type I error, which is the opposite of its performance in testing a binary trait.

For testing (7) the interaction in matched-pair data, the strategy of performing unrestricted permutation yielded extremely conservative type I error. Figure 7 confirmed the lack of power with unrestricted permutation.

The power and sensitivity of various 1:*m* matched study designs were contrasted in Figures 8–10. Figure 8 showed that the matched-pair design always gained substantial efficiency over an analysis of data from an equivalent number of *independent* cases and controls over a wide range of *θ*_2_ values. Figure 9 shows that, with a fixed number of total samples, maximizing the number of distinct sets (i.e., using 1:1 pairs) rather than increasing the number of controls per set optimized efficiency. In Figure 10, we show that adding more control samples to each set, while keeping the number of sets fixed, has a relatively small effect on power and sensitivity; the addition of each successive control sample yielded diminishing returns. Taken together, Figures 8-10 suggest that when data have a matched structure, a matched analysis outperforms an unmatched analysis and, in general, increasing the number of controls in a 1:*m* matched study beyond 1:2 may only result in fairly small improvements in power and sensitivity.

**Fig 8.**
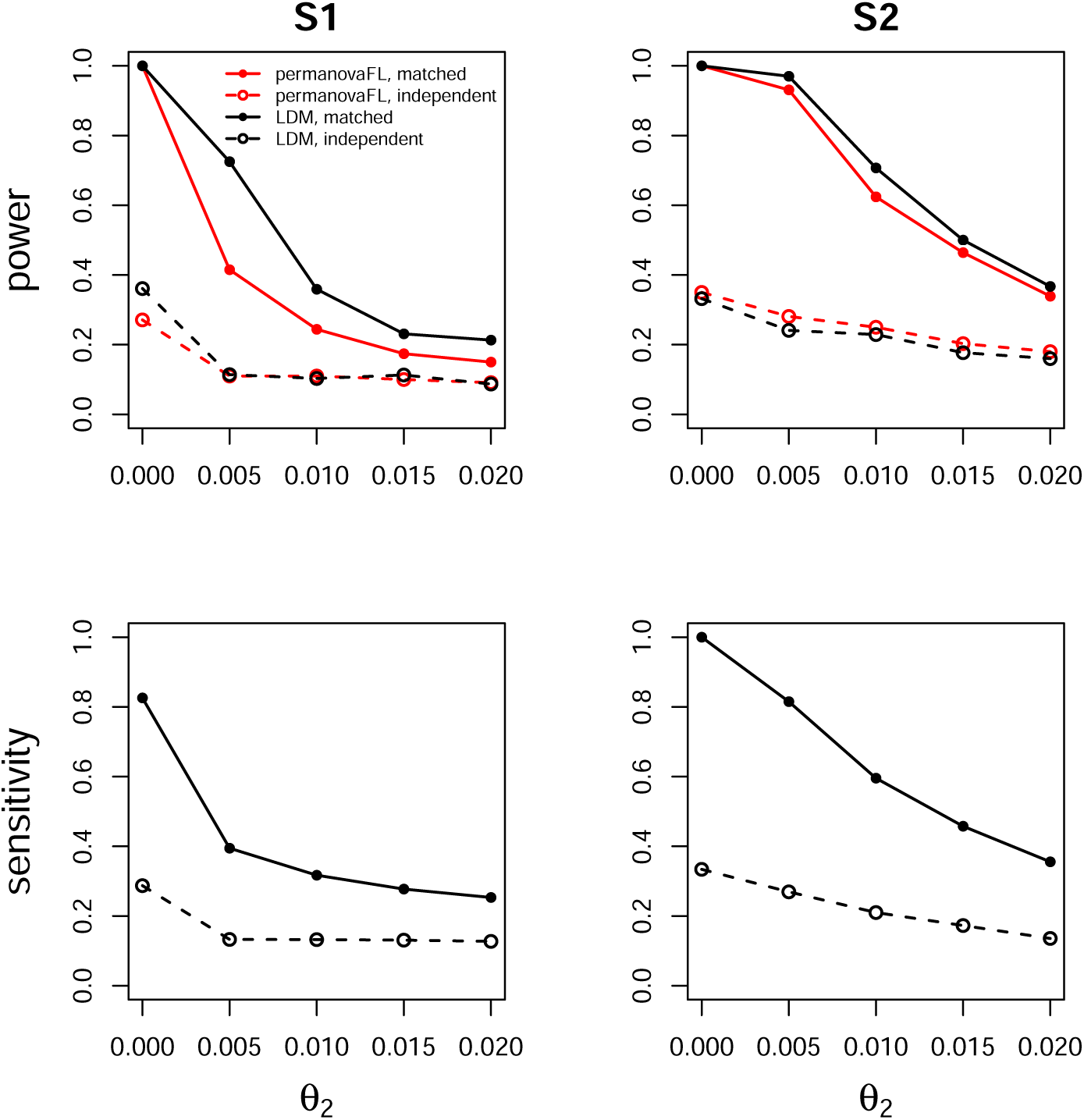
Comparing the matched-pair design (solid lines) with the independent case-control design (dashed lines) over varying sample-level heterogeneity *θ*_2_. The effect size *β* was set to 0.1 (S1, power), 0.25 (S2, power), 0.8 (S1, sensitivity), and 0.6 (S2, sensitivity). All simulations shown here use between-set heterogeneity parameter *θ*_1_ = 0.02.

**Fig 9.**
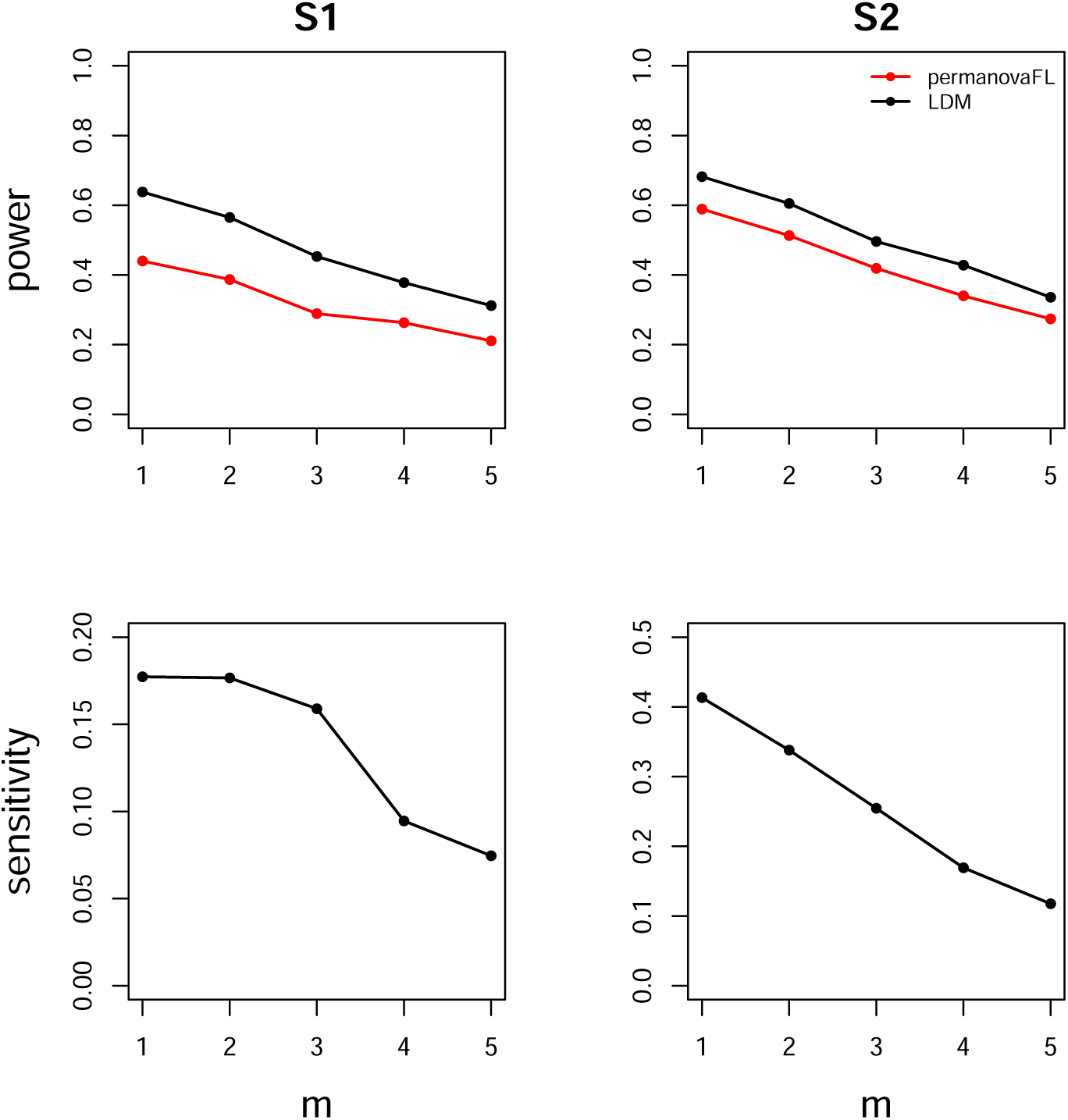
Comparing various 1:*m* matched-set designs with a fixed number (90) of total samples. The effect size *β* was set to 0.12 (S1, power), 0.22 (S2, power), 0.5 (S1, sensitivity), and 0.46 (S2, sensitivity). All simulations shown here use between-set heterogeneity parameter *θ*_1_ = 0.02 and within-set heterogeneity parameter *θ*_2_ = 0.007.

**Fig 10.**
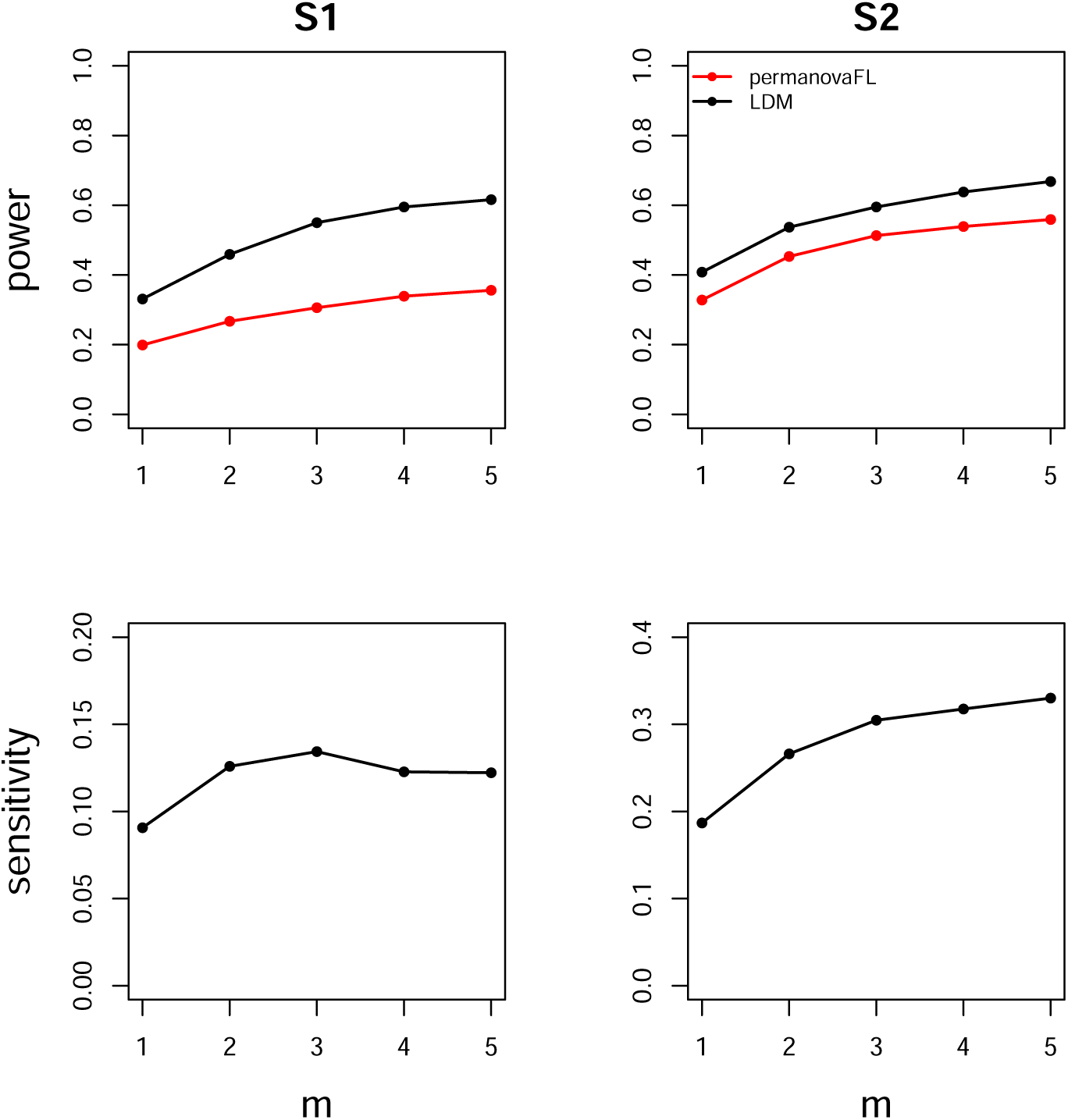
Comparing various 1:*m* matched-set designs with a fixed number (50) of total sets. The effect size *β* was set to 0.08 (S1, power), 0.16 (S2, power), 0.35 (S1, sensitivity), and 0.34 (S2, sensitivity). All simulations shown here use between-set heterogeneity parameter *θ*_1_ = 0.02 and within-set heterogeneity parameter *θ*_2_ = 0.007.

### Analysis of the MsFLASH data

The data for our first example were extracted from the study “Menopause Strategies: Finding Lasting Answers for Symptoms and Health” (MsFLASH) [9, 10]. This double-blinded, randomized trial enrolled women into one of three-arms: oral estradiol (arm 1), oral venlafaxine (arm 2) (two commonly used drugs to alleviate menopausal hot flashes) or placebo (arm 3). To examine the effect of these drugs on the vaginal microbiome, 113 vaginal swab samples were collected at baseline (before treatment), and at weeks 4 and 8 post-treatment. 16S rRNA gene sequencing was performed, and the results were summarized into 171 OTUs. Specifically, 9 sets (women) in the estradiol arm, 10 in the venlafaxine arm, and 18 in the placebo arm have data from swab samples at all three visits; one woman in the estradiol arm only provided samples at baseline and week 4. Due to the small sample size, we also considered an enlarged “treatment” group that combined the estradiol and venlafaxine arms. The ordination plot (Figure 11) showed that the samples from the same woman tended to cluster together.

**Fig 11.**
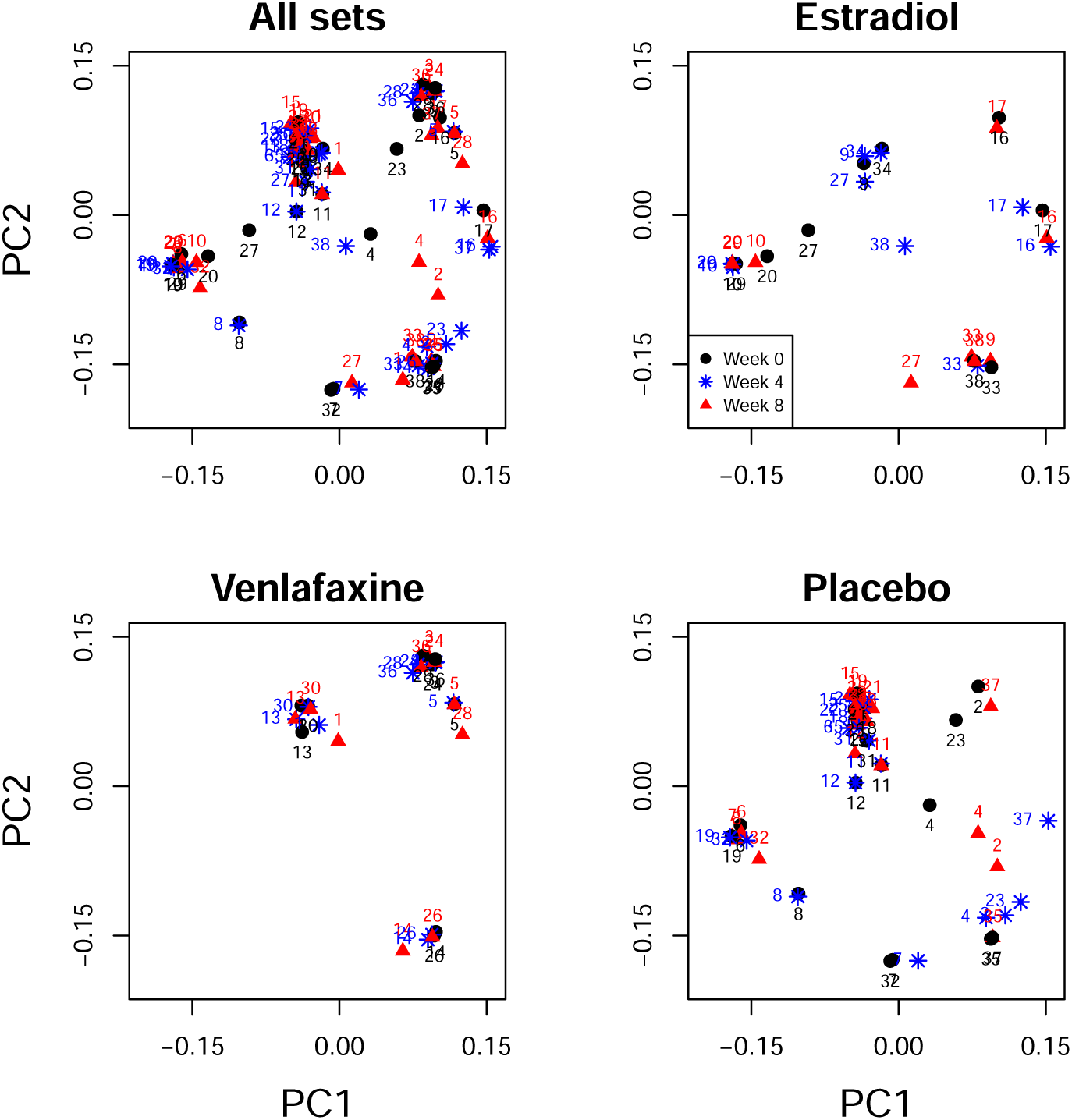
Ordination plots for the MsFLASH data. The texts above the symbols are the set IDs. The plot entitled “All sets” show the original ordination. The three plots entitled “Estradiol”, “Venlafaxine”, and “Placebo” show the stratified ordination by the three arms for the sake of clarity (using the same coordinates as in the plot entitled “All sets”).

In each arm, we tested whether the composition of the vaginal microbiome changed between baseline and week 4, baseline and week 8, and weeks 4 and 8; each of these tests was based on 1:1 paired data. We also tested the microbiome differences pre- and post-treatment by comparing baseline and post-treatment (both week 4 and week 8) samples without differentiating between time since treatment using a 1:2 matched-set design; the estradiol arm was an exception, as one set had only two samples, resulting in unbalanced data. We applied the LDM (using the omnibus test) and `permanovaFL` (using the Bray-Curtis distance) with the proposed strategy. As a comparison, we also applied DESeq2 (adjusting for the set ID) and the Wilcoxon signed-rank test to 1:1 matched data.

We limited our analysis for each arm to OTUs that were present at least 5 times in each of the four subsets of samples, which resulted in, for example, 31 OTUs in the venlafaxine arm. All results were summarized in Table 2. Only the comparisons within the venlafaxine arm yielded some significant *p*-values (*<* 0.05). In particular, the LDM generated *p*-value 0.033 for comparing the baseline and week 4 samples, followed by a smaller *p*-value 0.0042 for the baseline and week 8 samples, and then the smallest *p*-value 0.0003 for the baseline and the combined week 4 and week 8 samples. These *p*-values suggested an effect of venlafaxine on the vaginal microbiome, which was strengthened at week 8 relative to week 4. However, the differences between week 4 and week 8 were not found to be significant (the LDM *p*-value = 0.76). The results of `permanovaFL` corroborated these conclusions. In the comparison of the baseline vs. weeks 4 and 8, the LDM detected four OTUs (*Campylobacter, Gardnerella vaginalis, Porphyromonas*, and *Aerococcus christensenii*) to be differentially abundant at the nominal FDR 20% (we chose a relatively high nominal FDR because of the small number of sets), whereas DESeq2 detected none and the Wilcoxon test was not applicable.

**Table 2.**
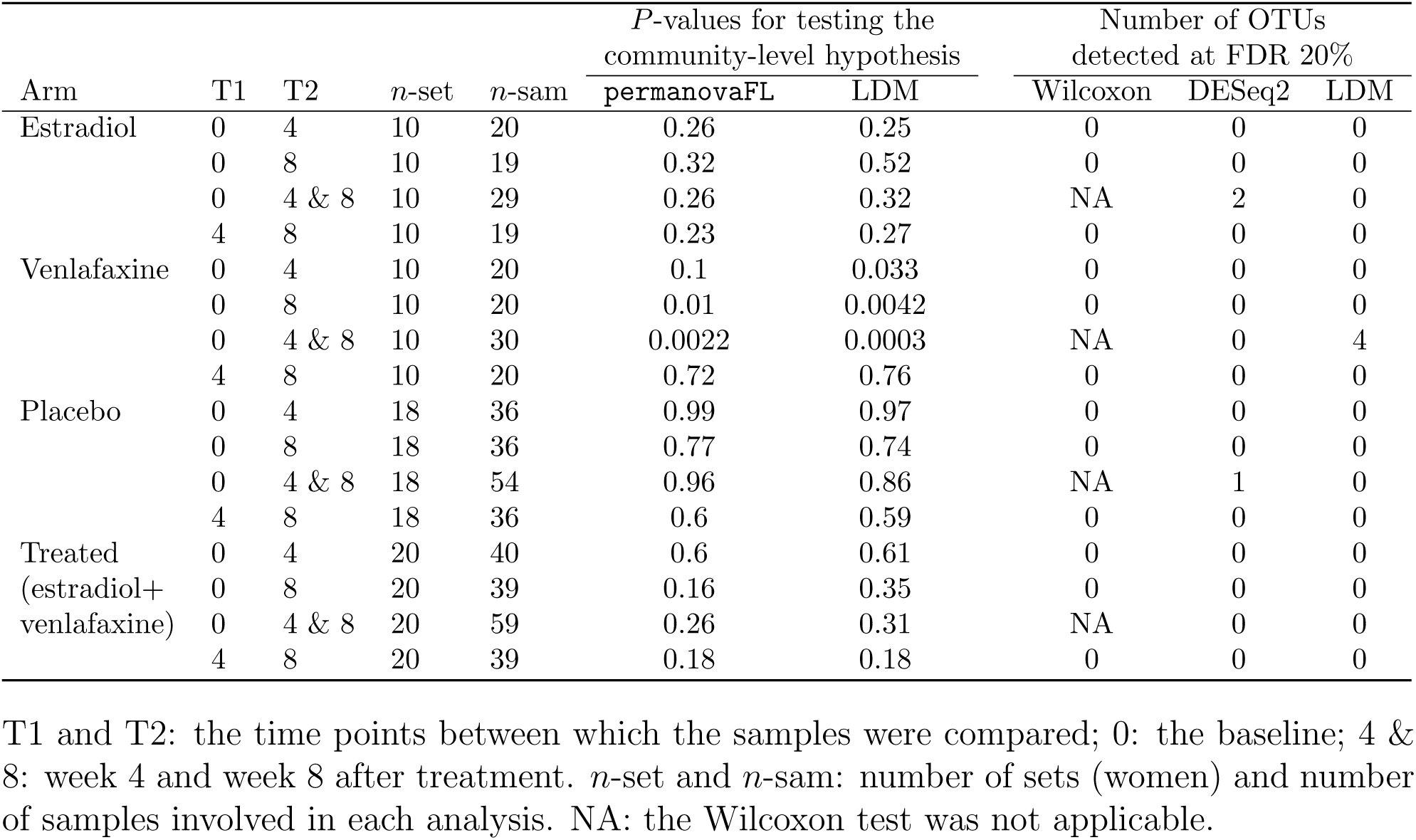
Results in analysis of the MsFLASH data.

| Arm | T1 | T2 | *n*-set | *n*-sam | *P*-values for testing the<br>community-level hypothesis |  | Number of OTUs<br>detected at FDR 20% |  |  |
| --- | --- | --- | --- | --- | --- | --- | --- | --- | --- |
|  |  |  |  |  | permanovaFL | LDM | Wilcoxon | DESeq2 | LDM |
| Estradiol | 0 | 4 | 10 | 20 | 0.26 | 0.25 | 0 | 0 | 0 |
|  | 0 | 8 | 10 | 19 | 0.32 | 0.52 | 0 | 0 | 0 |
|  | 0 | 4 & 8 | 10 | 29 | 0.26 | 0.32 | NA | 2 | 0 |
|  | 4 | 8 | 10 | 19 | 0.23 | 0.27 | 0 | 0 | 0 |
| Venlafaxine | 0 | 4 | 10 | 20 | 0.1 | 0.033 | 0 | 0 | 0 |
|  | 0 | 8 | 10 | 20 | 0.01 | 0.0042 | 0 | 0 | 0 |
|  | 0 | 4 & 8 | 10 | 30 | 0.0022 | 0.0003 | NA | 0 | 4 |
|  | 4 | 8 | 10 | 20 | 0.72 | 0.76 | 0 | 0 | 0 |
| Placebo | 0 | 4 | 18 | 36 | 0.99 | 0.97 | 0 | 0 | 0 |
|  | 0 | 8 | 18 | 36 | 0.77 | 0.74 | 0 | 0 | 0 |
|  | 0 | 4 & 8 | 18 | 54 | 0.96 | 0.86 | NA | 1 | 0 |
|  | 4 | 8 | 18 | 36 | 0.6 | 0.59 | 0 | 0 | 0 |
| Treated<br>(estradiol+<br>venlafaxine) | 0 | 4 | 20 | 40 | 0.6 | 0.61 | 0 | 0 | 0 |
|  | 0 | 8 | 20 | 39 | 0.16 | 0.35 | 0 | 0 | 0 |
|  | 0 | 4 & 8 | 20 | 59 | 0.26 | 0.31 | NA | 0 | 0 |
|  | 4 | 8 | 20 | 39 | 0.18 | 0.18 | 0 | 0 | 0 |

T1 and T2: the time points between which the samples were compared; 0: the baseline; 4 & 8: week 4 and week 8 after treatment. *n*-set and *n*-sam: number of sets (women) and number of samples involved in each analysis. NA: the Wilcoxon test was not applicable.

Motivated by the likely trend of strengthened effect of venlafaxine over time, we reanalyzed the data at weeks 0, 4, and 8 in the venlafaxine arm, treating “week” as a quantitative variable. However, this analysis yielded *less* significant global *p*-values (0.043 by the LDM and 0.096 by `permanovaFL`), suggesting that the change in OTU frequencies as a function of time since treatment initiation is probably non-linear. We also tested whether the effect of venlafaxine is the same for the five white and four black women (excluding one women in the “other” race category), i.e., we tested the interaction between week (coded as 0 vs. 4 & 8) and race. The global *p*-values are 0.44 by the LDM and 0.39 by `permanovaFL`, suggesting no racial difference in the effect of vanlafaxine.

### Analysis of the Alzheimer’s disease data

The data for our second example were generated from a pair-matched study comparing the gut microbiome of 25 patients with Alzheimer’s disease (AD) and their age- and sex-matched controls [11]. A covariate of particular interest was the APOE *ϵ*4 genotype, which was coded as carriers (one or two *ϵ*4 alleles) vs. non-carriers (zero *ϵ*4 alleles). APOE *ϵ*4 genotype is a potential confounder of the association between the gut microbiome and AD, as it is distributed differently in the AD patients than in the controls (AD: 72% carriers; control: 20% carriers; *p*-value*<*0.001) in the study sample, and has been found to influence the gut microbiome [12]. Since matching on APOE *ϵ*4 genotype was not used in the study design, it should be adjusted for in the association test. The microbiome data were summarized into 972 OTUs, of which 723 were present at least 5 times in the study sample and included in our analysis. We applied the same methods as those for the MsFLASH data, except we do not report results for the Wilcoxon signed-rank test, which is not applicable in the presence of a within-pair covariate.

The results were summarized in Table 3. Without adjustment of the APOE *ϵ*4 genotype, the LDM yielded *p*-value 0.0001 for testing the community-level association and detected 66 OTUs (at nominal FDR 10%) that were differentially abundant between AD patients and controls. After adjustment for APOE *ϵ*4 genotype, the LDM yielded *p*-value 0.0159 and detected no OTUs. The results of `permanovaFL` corroborated this conclusion. These results suggest that much of the association seen without adjusting for APOE *ϵ*4 genotype is due to confounding.

**Table 3.** Results in analysis of the Alzheimer’s disease (AD) data

|  | *P*-value for testing the<br>community-level hypothesis |  | Number of OTUs<br>detected at FDR 10% |  |
| --- | --- | --- | --- | --- |
|  | permanovaFL | LDM | DESeq2 | LDM |
| Without adjustment of APOE | 0.0001 | 0.0001 | 168 | 66 |
| With adjustment of APOE | 0.0069 | 0.0159 | 66 | 0 |

## Discussion

We have developed a novel strategy that extends PERMANOVA (implemented in both `adonis2` and `permanovaFL`) and the LDM for analyzing matched-set microbiome data, that can account for complex design features such as unbalanced data, sample-level confounding covariates, and continuous traits of interest. This strategy corresponds to a specific application of PERMANOVA and the LDM, without modifying any of their methodologies. Our simulations show that the proposed strategy was the most efficient among all strategies we considered, when applied to either PERMANOVA or the LDM. The LDM was also superior to existing methods, such as DESeq2 and the Wilcoxon signed-rank test, for testing individual OTUs with matched-set data. In addition, our simulation studies suggested that the 1:1 matched-pair study is the most efficient design as it maintains a good balance between sequencing cost and statistical power.

Our results in analysis of the MsFLASH data did not agree with those reported by Zhao et al. [2], who found significant effects in the “treated” group only (rather than the venlafaxine arm). Their method was based on log-ratio-transformed frequency data and used a pseudo count value of 0.01 for zero count data, which essentially resulted in a different hypothesis being tested than that used in our methods. Similarly, we found that much of the association reported by [11] between AD disease status and the microbiome may be due to confounding by APOE *ϵ*4 genotype. This finding emphasizes the need to develop and use microbiome methods, such as those we have reported here, that can account for complex design features, like matching with within-set confounding covariates, that are often found in epidemiological studies involving the microbiome.

Hu and Satten [5] have shown that for independent case-control samples, the power of the LDM was sensitive to the OTU data scale, i.e. if untransformed frequency scale or arcsin-root-transformed data were used. We found (Figure S1) that these patterns persisted in the analysis of matched-set data. As a result, we reiterate the recommendation in [5] and use the *omnibus* test for the LDM, which corresponds to the minimum of the *p*-values obtained on the frequency and arcsin-root-transformed scales.

The strategy we have proposed here is applicable to any matched-set microbiome data as long as model residuals can be assumed to have an exchangeable correlation structure. In some settings, longitudinal microbiome data that have time-varying traits (i.e., time, antibiotic intake, or infection) can be reasonably assumed to have an exchangeable correlation structure. The simple within-cluster permutation approach used here is not valid for other correlation structures such as the autoregressive model. We are currently developing methods for analysis of clustered or longitudinal microbiome data having an arbitrary residual correlation structure.

Our simulation studies showed that matched-set sampling, when available, can result in a substantial increase in power to detect global associations and sensitivity to detect individual OTUs when our approach is used. This is presumably because the overdispersion parameter for the matched data is smaller than it is for independent data sampled from the same population. In the independent data sample, the overdispersion parameter describing each observation is effectively the sum of the between- and within-set heterogeneity parameters (*θ*_1_ and *θ*_2_ in our simulations). In the matched data, the between-set heterogeneity (represented by *θ*_1_ in our simulations) is effectively conditioned out. Thus, we expect the advantage of a matched analysis over an unmatched analysis to increase as the between-set heterogeneity increases. Presumably when the within-set heterogeneity is large compared to the between-set heterogeneity, a matched analysis would have a smaller advantage.

## Disclaimer

The findings and conclusions in this report are those of the authors and do not necessarily represent the official position of the Centers for Disease Control and Prevention.

## Acknowledgement

The authors would like to acknowledge the Wisconsin ADRC’s biostatistical support provided by the Data Management and Biostatistics Core.

## Availability of data and materials

The R package `LDM` is available on GitHub at https://github.com/yijuanhu/LDM in formats appropriate for Macintosh, Linux, or Windows systems.

## Authors’ contributions

YJH conceived the study, developed the method, performed simulation studies, analyzed the data, and wrote the manuscript. GAS conceived the study, developed the method, and wrote the manuscript. ZZ performed simulation studies and analyzed the data. CM provided the biological insights and interpretation for the results from analyzing the MsFLASH data. All authors read and approved the final manuscript.

## Consent for publications

## Ethics approval and consent to participate

This study only involved secondary analyses of existing, de-identified datasets; as such it does not require separate IRB consent.

