## Supplemental Figure S1 for "Analyzing matched sets of microbiome data using the LDM and PERMANOVA"

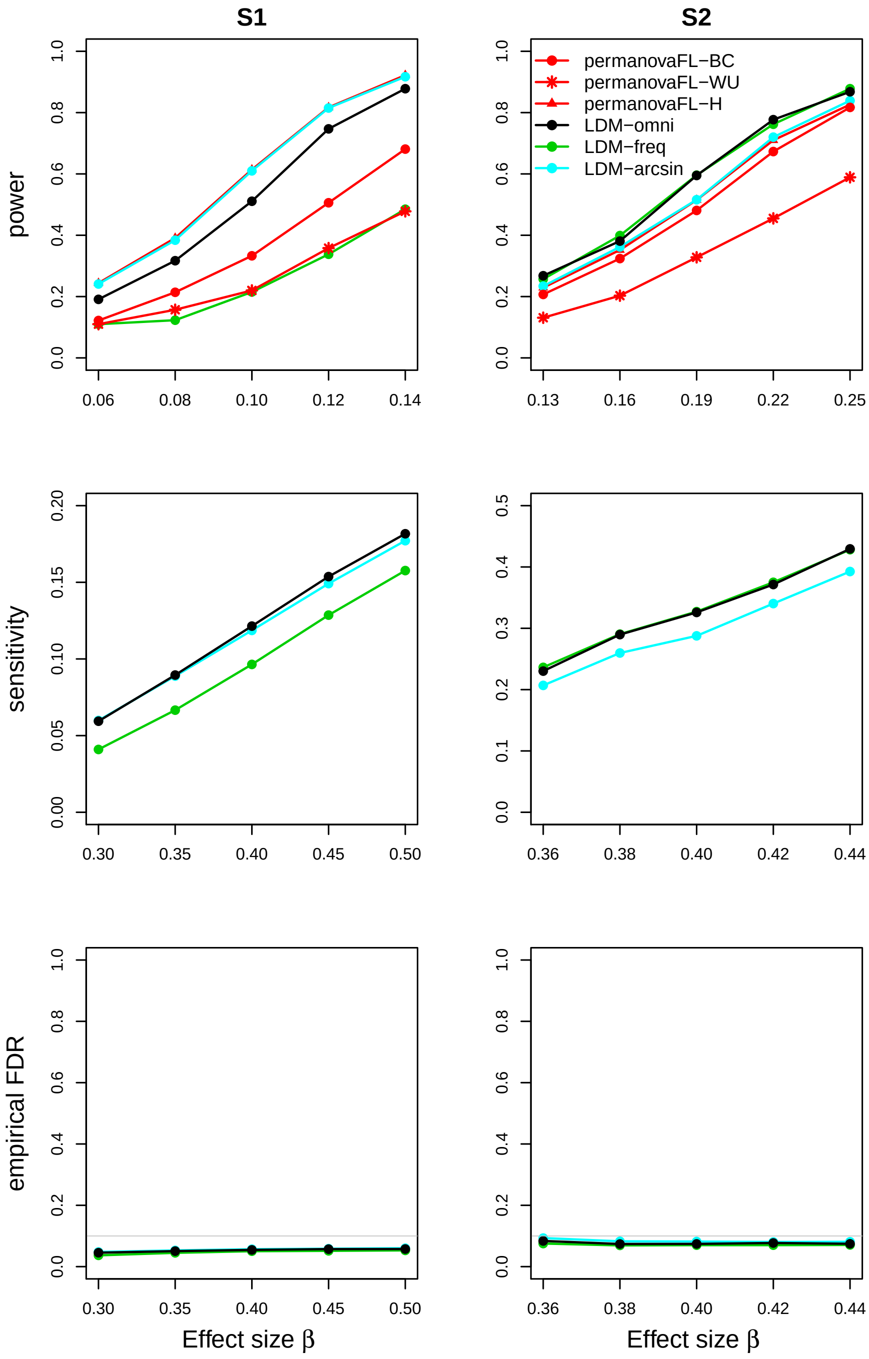


**Fig S1**. Simulation results for the matched-pair data of scenario (1). The LDM on frequency scale (freq), arcsin-root transformed frequency scale (arcsin), and the omnibus test (omni) are shown. `permanovaFL` based on the Bray-Curtis (BC), weighted UniFrac (WU), and Hellinger (H) distances are shown. All methods adopted the proposed strategy.
